# Mechanics of *Hydra* Detachment from Substrates: The Role of Substrate Rigidity and Starvation

**DOI:** 10.1101/160028

**Authors:** Neha Khetan, Shagun Maheshwari, Chaitanya A. Athale

## Abstract

*Hydra* is a fresh water hydrozoan living as a solitary polyp with a sedentary feeder lifestyle attached to a substrate. In times of food shortage they are reported to detach from their substrate and move either by drifting or ‘somer-saulting’. The attachment to the substrate is usually by the basal-body which secretes a mucosal adhesive. The mechanical strength of the adhesion of Hydra has not been quantified so far. Here, we measure the force required to detach *Hydra vulgaris* and *Hydra magnipapillata* from a surface and the role of physical and physiological factors. In order to do this, we have developed a flow chamber with a calibrated jet of water. We find *H. vulgaris* adhering to a hard substrate - a glass cover slip- requires more force to detach it as compared to a soft substrate- polyacrylamide gel. While *H. vulgaris* after one week of starvation detaches with very similar values of stress, *H. magnipapillata* detaches more readily when starved. These results suggest that the strength of adhesion is strongly affected by the stiffness of the substrate, while nutritional status dependence of detachment force appears to be species dependent. Given that *Hydra* detachment is required during locomotion, our measurements on the one hand suggest the magnitude of forces the animal must exert to detach itself. Additionally, our results suggest active detachment of the base might be required for *Hydra* to achieve movement, and only a small contribution coming from weakening adhesion.

## 1 Introduction

Aquatic life forms ranging from single-celled to multi-cellular have evolved a variety of strategies to remain static through adhesion to substrates. The specific mechanism by which they achieve this adhesion ranges from suckers and nanometer scale spatulae to biological adhesives (reviewed by Gorb (2008)). Amongst aquatic animals the adhesion of the mussel *Mytilus edulis* has been particularly well studied (reviewed by (Waite, 2002)). The mussel shells attach to rocky substrates with byssal threads with multiple proteins contributing differing mechanical properties (Lin et al., 2007), of which the amino acid 3,4-dihydroxy-L-phenylalanine (dopa) is considered a vital component (Lee et al., 2006).

*Hydra* on the other hand, are fresh water dwelling Hydrozoans of phylum Cnidaria that live as solitary polyps, typically found attached to substrates like stems, branches or leaves under-water. Renewed interest in *Hydra* is due to its regenerative ability, along with genome sequence and the evolutionary relatedness of the regenerative pathways to vertebrates (Fujisawa, 2006; Watanabe et al., 2009). In their natural environment, *Hydra* are subject to gentle flows and so far their movement has been attributed to passive drifting. Wagner has noted that the resistance of an attached *Hydra* to water flows might be an adaptation to the diverse environmental conditions (still and flowing water) that it is exposed to and its inability to actively swim, once suspended in water (Wagner, 1905). The movement of *Hydra* that are already attached to a substrate was observed to occur by ‘somersaulting’-animals attach their tentacles to a new position, detach the basal disk with body contraction, straighten and reattach the basal disk at a new position near the hypostome (Wagner, 1905). Annandale reported *Hydra vulgaris* from Indian samples to be observed to be actively ‘crawling’, which involved similar ‘somersaulting’ motion (Annandale, 1911). The movement was thought to enable the individual to leave unfavourable environments. While it is known that muscles that are ectodermal and longitudinal drive contraction, while endodermal circular muscles drive extension of *Hydra*, the biomechanics of *Hydra* detachment has yet to be examined.

In its sedentary mode *Hydra sp.* is attached by its basal-body also called the basal disk or ‘foot’ to substrates by a glandular secretion (Brien, 1960). Like other parts of Hydra, the animal can also regenerate the basal disk when amputated (Amimoto et al., 2006). Histologically the cells of the disk consist of the inner endoderm and outer ectoderm. The cells are glandular, conical in shape and filled with granules (Bode et al., 1986). The cells secrete large amounts of mucus needed for the attachment of the animal to substrates. The basal disk was thought for long to be a closed structure but more recently a pore-like structure in the disk called the aboral pore (Shimizu et al., 2007) has been found. While the histology of the foot is understood, the nature of the mucus as a bioadhesive which works under water could be interesting both from a fundamental perspective of adhesives, as well as applications in biocompatible materials (Waite, 2002). Recently, the glue from *Hydra magnipapillata* has been characterised and found to be based on gylcans and glycoproteins (Rodrigues et al., 2016a). Additional gene-expression analysis has revealed 21 transcripts to be expressed in the basal disk alone (Rodrigues et al., 2016b). While remaining attached is important for *Hydra*, the ability to detach and move is equally important. It would appear addressing the biomechanics of detachment, could connect the mechanics of muscle-generated forces with the biochemistry of ‘glue’ attachment.

Measuring the force for detachment of larger animals such as *Mytius sp.* has involved mechanical spring-based instruments (Bell and Gosline, 1996; Denny, 1987), while sea anemone detachment has been measured using force transducers (Koehl, 1977). Such instruments however do not mimic the naturally occurring flows that aquatic animals are likely to experience and measurements could suffer from artefacts from mechanical contact. Flow chambers address some of these shortcomings and have been used to study biofouling using turbulent (Schultz et al., 2000) or pumped flows coupled to inline flow meters (Neal et al., 1996). Flow tanks that have been described for whole organism studies (Vogel and LaBarbera, 1978) and parallel-plate flow chambers used to studying leukocyte adhesion (Chen and Springer, 1999) are successful means to measure detachment dynamics of samples ranging from cells to whole organisms.

We have chosen to characterise the biomechanics of *Hydra sp.* detachment with calibrated fluid flows, from which we estimate the drag force required to displace the animal from a substrate. We use this device to measure the flow rate required to detach the *Hydra* from substrates of different stiffness and proceed to examine the role that starvation and substrate stiffness plays in the stress required to detach the individuals.

## 2 Results

### 2.1 *Hydra* sizes

In order to estimate the forces exerted by flow, we needed to morphologically characterise the *Hydra* we used in the study. Two species were chosen due to the differences in sizes and availability, namely *H. vulgaris* and *H. magnipapillata.* While qualitatively in a dissection microscope *H. vulgaris* was seen to be shorter than *H. magnipapillata*, their widths appeared comparable (Figure 1). This was confirmed by the estimate of the mean cross-sectional diameter of the foot of *Hydra vulgaris* to be 0.279 mm for (Figure 1(a)–(d)) and 0.342 mm for *H. magnipapillata* (Figure 1(e)–(h)). Given the base of the hydra is approximately circular, we estimated the mean base area of *H. vulgaris* and *H. magnipapillata* to be 0.24 and 0.37 *mm*^2^ respectively. These values show small differences in base-areas, while the length of *H. magnipapillata* is approximately two-fold greater than *H. vulgaris*, varying between individuals. For our measurements, the flow was directed to the base of the anim and is not expected to influence our measurement as a result. As a next step, we needed to calibrate the flow flow chamber.

**Figure 1:**
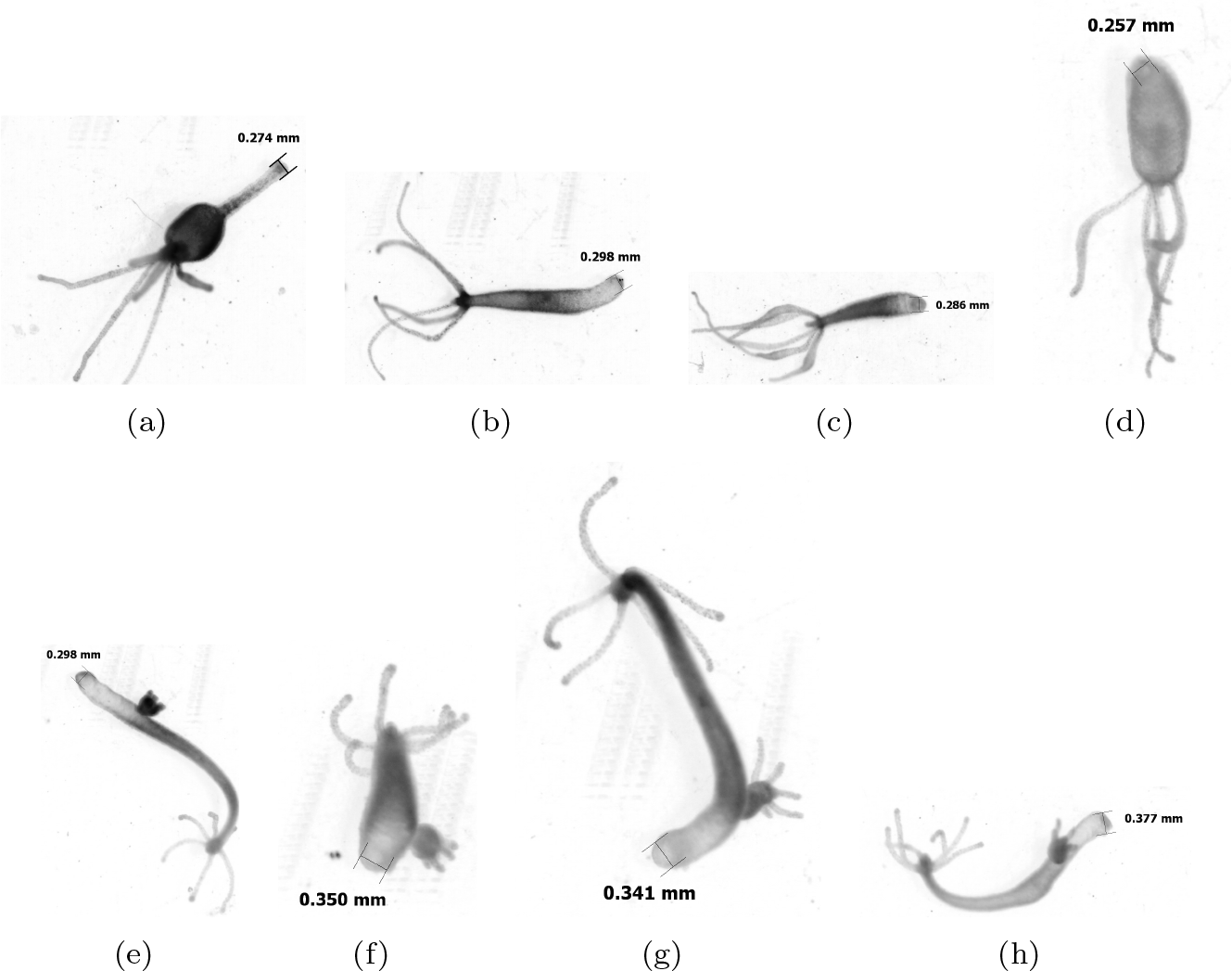
Estimating the size of *Hydra:* Images of live animals of (a)-(d) *Hydra vulgaris* and (e)-(h) *Hydra magnipapillata* under a dissection microscope in a drop of “M” solution. The scale bar in every image indicates the base diameter.

### 2.2 Characterizing the laminar range of the flow-chamber

The flow chamber setup consists of a syringe pump connected by tubing to a trough filled with medium, in which *Hydra* is submerged and subjected to the flows (Figure 2(a)). In order to characterise the nature of the fluid flow in the chamber, we needed to ensure the forces are generated due to laminar flows. Flowing safranin-stained water into the apparatus in the absence of any obstacle, turbulence was observed beyond the pipe exit (E) at a certain point of turbulence (T) onset (Figure 2(b)). This distance from the pipe exit (*E*) at which turbulence (*T*) due to eddies is observed was measured for multiple flow rates. The plot of distance (T) as a function of flow rate (Q) shows that even for the fastest flow-rates, the eddies begin 10 mm from the pipe exit (E) (Figure 2(c)). As a result, we placed the *Hydra* at 5 mm from E in subsequent experiments, to ensure that the force experienced is due to laminar flows alone.

In order to independently confirm the consistency of this data, we also estimated the Reynolds number for each as a function of flow rate (Q) using:

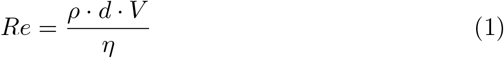

**Figure 2:**
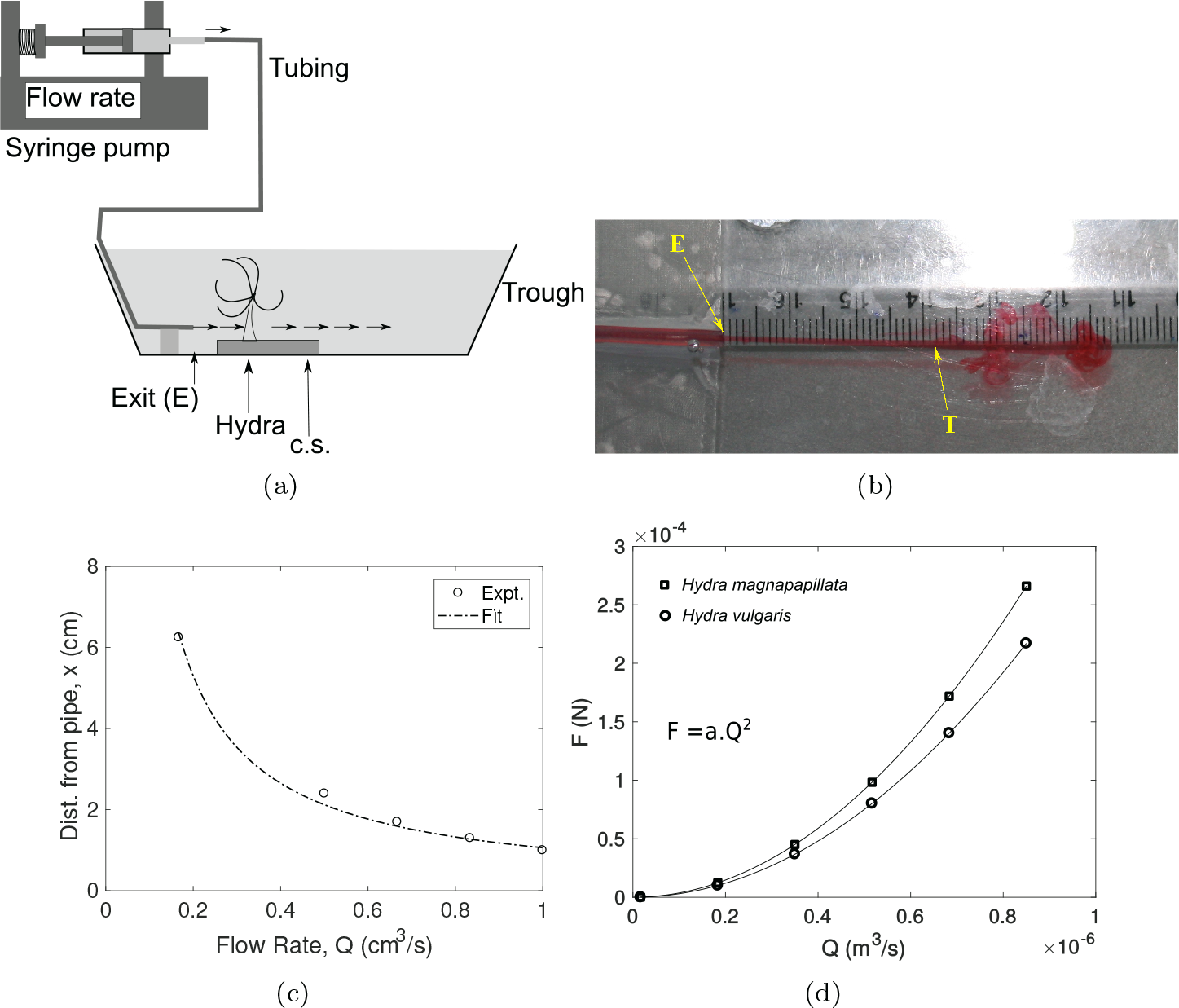
Detachment of Hydra by laminar flow set up. (a) A schematic representation of the experimental set up with the *Hydra* placed on a coverslip (c.s.) at a fixed distance from the pipe exit (E) submerged in buffer in a glass trough. The arrows indicate the direction of flow of water from the syringe pump. (b) The view from the top of the distance of the onset of turbulence (T) flow from the pipe exit (E) estimated by flowing safranin containing water. (c) The distance from E at which turbulence is seen is plotted against the flow rate (circle) and fit (dashed line) by the equation x = *c*_1_/*Q* (Equation 2). (d) The drag force experienced by Hydra due to flow is calculated (Equation 4) and fit by Equation 7. The fit parameter for *Hydra vulgaris* (circle) is *c*_2_ = 3 . 10^8^ kg .m^-5^ and for *Hydra magnipapillata* (square) is c_2_ = 3.68 . 10^8^ kg . m^-5^.

where, *Re* is Reynolds Number, ρ (*kg . m^-3^*) is the density of fluid, *d* (m) is the diameter of the pipe *V* (*m*. *s*^-1^) is the fluid velocity and η (*N*. *s*. *m*^-2^) is the dynamic viscosity. In order to estimate Re in terms of Q, we used the relation Q = V × A, where *Q* (*m*^3^. *s*^-1^) is the volume rate of the fluid and *A*(*m*^2^) is the area of cross section of the jet of fluid. The resulting values of Re (Table 1) appeared to converge on a mean value of Re = 1.3243(±0.8785).10^3^.

**Table 1:** Flow is laminar within the experimental range of flow rates.

| Flow rate, Q (ml/min) | Reynolds Number, Re |
| --- | --- |
| 10 | 264.863 |
| 20 | 529.726 |
| 30 | 794.589 |
| 40 | 1059.45 |
| 50 | 1324.32 |
| 60 | 1589.18 |
| 70 | 1854.04 |
| 80 | 2118.9 |
| 90 | 2383.77 |
| 100 | 2648.63 |

In order to relate the onset of turbulence with the distance from the pipe exit (x), we rewrite Equation 1 in the simpe form by using the relation V = Q/A and substituting all constant values in one lumped constant c_1_ as follows:

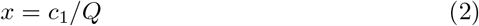

This equation is qualitatively comparable to the experimental estimates of x as a function of Q, i.e. an inverse relation (Figure 2(c)). The constant c is given by:

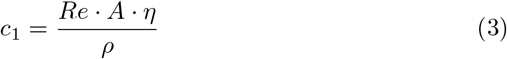

We substitute known values into this Equation 3 and estimate *c*_1_ to be 1.13 *cm*^4^/*s*, given: η = 1.002 . 10^-3^ *N*. *s/m*^2^ and ρ = 10^3^ *kg/m*^3^ and the cross-sectional area of the flow *a_f_* = 5.0265 . 10^-7^*m*^2^, since the jet diameter = 0.8 mm. When we compare the fit of Equation 2 to our data of turbulence onset distance (T) with Q (Figure 2(c)), the parameter 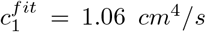 This validates our approach from first principles.

In addition, the onset of turbulence over the entire range of flow rates remains x≥ 1 cm. Thus, we confirm that the force experienced by objects at a distance of less than ∼ 1 cm can be treated as being due to a laminar flow for all values of Q.

### 2.3 Drag force experienced by *Hydra*

In order to estimate the force at which *Hydra* detaches from the substrate, we need to relate the volume flow rate with force. To this end, the drag-force (*F*_*drag*_) exerted by the fluid flow on *Hydra* was estimated from the drag-equation:

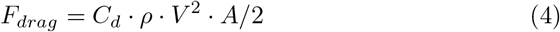

where *C*_*d*_ is the drag coefficient, *ρ* is the density of the fluid, V is the velocity of flow and A is the projected area of the body in the path of fluid flow. We can relate the linear velocity (*V*) of the fluid with the volume flow rate (Q) as follows:

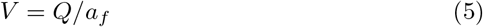

where *a_f_* is the cross-sectional area of the flow. On substituting V (Equation 5) in the expression for drag force (Equation 4) gives us:

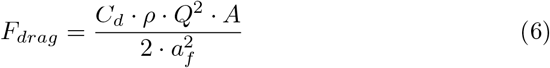

Thus the detachment force of *Hydra* can be calculated using Equation 6. The drag coefficient (C_d_) is dimensionless constant and depends on properties of the object and fluid such as shape and Reynolds number. For our calculations, choice of C_d_ was made by approximating the shape of *Hydra* to a cylinder with it’s long axis normal to the direction of flow. Based on the length:width ratio of *Hydra*, the drag coefficient of 0.68 was chosen, based on standard results for a cylinder with length to the diameter ratio of 2:1 (Stoecker, 2004). For the sake of simplicity, we assumed the shape to be constant for the *Hydra* across all the conditions.

The calculated estimates of force with increasing flow rate was fit to a function of the form:

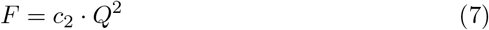

where Q is the volume flow rate (*cm*^3^/*s*) and *c*_2_ is a constant to compare the drag force dependence on flow rate between the species and the drag coefficients assumed.

### 2.4 Detachment force of *H. vulgaris* and *H. magnipapillata*

In order to measure detachment forces, individual *Hydra* were allowed to attach to a glass coverslip (coated or uncoated) in the incubator. At the time of the measurement, the coverslip was removed from the incubator and placed at the same distance from the pipe exit (x= 0.5 cm). The flow rate (Q) was gradually increased until the *Hydra* detaches. Measurements were repeated for ∼ 10 individuals of *H. vulgaris* and *H. magnipapillata.* By starving one set of animals, we addressed the effect of nutritional state. We also compared the effect of changing substrate stiffness on *H. vulgaris*. For each experimental condition, the flow rate at which *Hydra* detached was used to calculate the force of detachment (Equation 7). Using the estimate of the area of the basal disk, we thus estimate the shear stress of detachment. H. *vulgaris* detaches from glass substrates over a wide range of shear stresses: 0.6 to 1.8 MPa, for starved and fed samples (Figure 3(a)). H. *magnipapillata* detachment was only recorded in a few cases, since in most of the cases, as they did not detach within the maximal limit of ≤ 60 ml/min of the instrument (Figure 3(a)). Fed and unfed animals of *H. vulgaris* show a very minor difference in detachment shear stresses. The mean shear stress of detachment from a soft substrate of 5% polyacrylamide is lower than that measured on glass. However, more measurements will be required for statistical significance. However the shear stress required to detach *H. magnipapillata* is higher than that for *H. vulgaris* in the same nutritional state (Student’s paired t-test with 95% confidence interval). We therefore find the detachment shear stress to be in the range of 10^3^ N/m^2^, which is two orders of magnitude smaller than in molluscs (Denny, 1987). This suggests the *Hydra* adhere to their substrate with a weak glue in a substrate-stiffness dependent manner.

**Figure 3:**
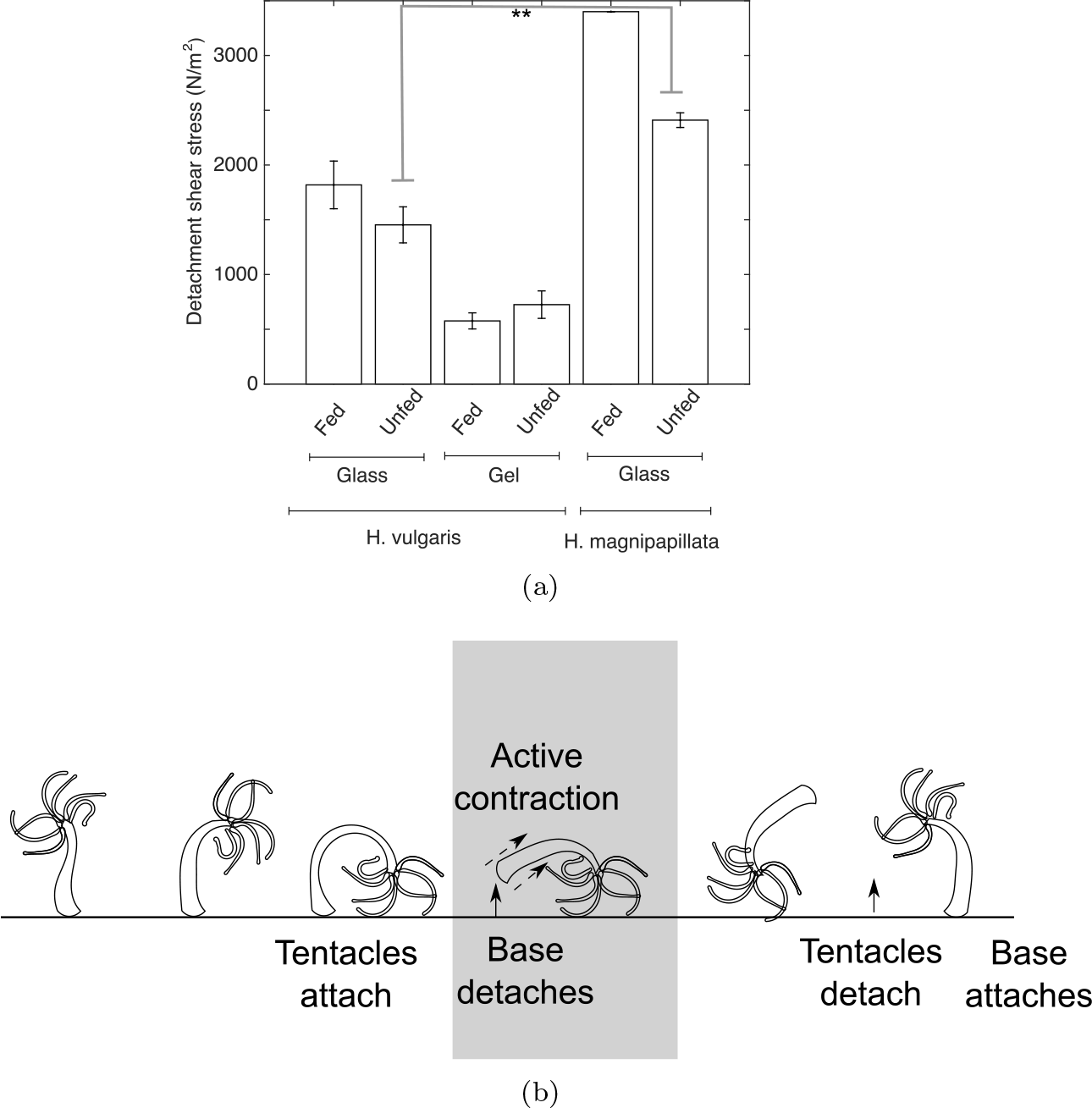
Detachment shear stress: (a) The mean shear stress (± S.E.) of detachment for *H. vulgaris* and *H. magnipapillata* under fed and unfed conditions for individual hydra attached to glass (stiffness 7.29 . 10^7^ kPa). *H. vulgaris* detachment was also tested on a 5% polyacrylamide gel (stiffness ∼ 8 kPa). The mean shear stress was compared using a pairwise, two-sided t-test with α = 0.05 (**). (b) The schematic depicts the role of active detachment of the base from the substrate during ‘somersaulting’ movement by *Hydra.* The detachment shear stress measurements presented here could suggest the amount of force required to be generated by active contraction of *Hydra.*

## 3 Conclusions and Discussion

Here for the first time we have quantitatively characterised the mechanics of *Hydra* detachment from an attached position on a substrate. We use a simple flow device which generates known amounts of shear stress through flows generated in the aqueous. We can show that smaller shear stresses are required to detach *H. vulgaris* from soft as compared to hard substrates. We find *H. vulgaris* and *H. magnipapillata* show little difference between fed and unfed (starved for one-week) conditions.

The steady attachment of sedentary aquatic animals can occur by multiple mechanisms, but the strength of the attachment to the substratum is related to its behaviour as well as the flows in which it lives. Typically free flowing streams with a gentle flow are reported to have flow speeds in the range of 0.5 m/s to 3 m/s. We can make an order of magnitude estimate of the shear stress (S) based on the fluid drag-force (*F_drag_*) from Equation 4 which simplifies to *S* = ρ.*v*^2^/2.*r* using the *C*_*d*_ of 0.68 based on the 2:1 ratio of length to the diameter (Stoecker, 2004) of *Hydra* and assuming the *Hydra* can be treated as cylinders, so the projected half-area of the a cylinder is affected by drag. To estimated the projected areas of *H. vulgaris* and *H. magnipapillata* the height is measured to be approximately 3 and 5 mm respectively and radius (r) is the same as in Section 2.1. As a result *H. vulgaris* will be expected to experience shear stresses between 1.8 × 10^3^ and 6.6 × 10^4^ N/m^2^, while *H. magnipapillata* is expected to experience between 2.5 × 10^3^ and 8.9 × 10^4^ N/m^2^. Given that we measure detachment shear stresses for both species ranging between 6 × 10^2^ to 3 × 10^3^ N/m^2^ (Figure 3(a)), it would suggest normal flows that the animal is likely to experience, would be sufficient to detach the animal from the substrate. This would suggest, that in addition to active motion, *Hydra* can also be passively detached from its substrate. This is corroborated by observations of drifting animals found in their natural habitat. Careful observations in still and moving streams combined with measurements of flow-rates *in situ* could be potentially used to test this prediction.

The measurements we report here are made on two kinds of artificial substrates-glass and 5% polyacrylamide gel. While glass is very stiff with a Young’s modulus of 72.9 MPa, the gel used has a reported stiffness of ∼8 kPa (Tse and Engler, 2010). Our measurements suggest the *Hydra vulgaris* is less firmly attached on a soft substrate as opposed to a hard substrate. The comparison with the bulk modulus of elasticity of freshwater aquatic plant leaves, which ranges between 1 and 10 MPa (Touchette et al., 2014), would suggest our measurements cover the range of stiffness that *Hydra* could be expected to encounter when attached to leaves. While on the one hand the differences in detachment are less than an order of magnitude and subject to large variations, it would be interesting in future to systematically vary substrate stiffness and examine the role it plays in movement of the animal. Additionally, the mechanical properties of the specific plants and other objects to which *Hydra* is naturally found attached to, could also determine whether there is any role at all for substrate stiffness.

In our experiments, we have used inert substrates during the detachment measurements, in order to study the role of mechanical properties in the absence of any material properties. However, it could have been possible that the chemical nature of the ‘glue’ might also be modulated during detachment, such as by hydrolysis by some enzyme produced by the animal itself. However a recent study that investigated the glue concluded that active attachment is more likely to be the primary method by which *Hydra* achieves detachment (Rodrigues et al., 2016a). While our data corroborates this by an independent means, since we do not observe a clear starvation dependent weakening of the bond, it would be useful in future to use motion-capture to carefully test these theories of ‘somersaulting’ movement of *Hydra* to capture the entire cycle of movement (Figure 3(b)) and the mechanics involved.

The force required to detach *Hydra* is two orders of magnitude smaller than the detachment stress of 1.2.10^5^ *N/m*^2^ reported for the well studied sedentary marine mussel *Mytilus sp.* (Denny, 1987). We hypothesise that the difference in habitat of *Hydra sp.* which mostly inhabits ponds and slow-flowing streams means that the detachment stresses do not need to be as high as those observed in *Mytilus* mussels, typically found attached to inter-tidal rocks subject to constant wave action (Bell and Gosline, 1996). Based on reports by Annandale and others, it is reasonable to assume this weaker adhesion of *Hydra* is an adaptation to the forces generated by currents it usually experiences in it’s natural habitat and for the mode of motility that it adopts.

The measurement setup, while simple, provides useful initial answers to a mechanical approach to animal behaviour. Potentially in future higher flow rate methods would require taking into consideration the turbulent regime (Schultz et al., 2000). In addition the starvation conditions in the native environment that trigger ‘somersaulting’ movement are not clearly reported. In our work we have empirically chosen a week of starvation. In future a controlled study on the factors and duration of nutrient withdrawal, combined with mechanics could help us better understand the triggers that govern the decision of *Hydra* to move.

In conclusion we have characterised the shear stress of detachment of two species of *Hydra* and find them to range between 0.14 to ≥ 0.8 kPa. We have shown for the species examined, the detachment stress is independent of the nutritional state (i.e. fed as compared to starved for one week) and only weakly dependent on substrate-stiffness. Additionally we find *H. magnipapillata* is detached at a higher stress value as compared to *H. vulgaris.* It leads us to hypothesise that the active detachment of *Hydra sp.* is likely to be driven by active muscle contractions. This work sets the stage for a more comprehensive study of the mechanics of locomotion by this organism.

## 4 Materials and Methods

### 4.1 Growth and handling of *Hydra*

*Hydra vulgaris Ind-Pune* (Figure 1(a)–(d)) and (Reddy et al., 2011) and *Hydra magnipapillata* (Figure 1(e)–(h)) were obtained from ARI (Pune, India). They were maintained in ∼ 200 ml of “M” solution containing 0.1 mM KCl, 1 mM *NaCl*, 1 mM *CaCl_2_. 2H_2_O*, 1 mM Tris (pH 8) and 0.1 mM *MgSO_4_. 7H_2_O* in water (Sugiyama and Fujisawa, 1977). The animals were maintained at 18 °*C* in an incubator with a lamp with timer kept on for ∼ 12h to artificially induce day-night cycles (Thermo Scientific, USA) and the beaker cleaned on a daily basis. *Hydra* were fed two to three hatched *Artemia sp.* (brine shrimp) every two days that were grown in a 0.6% saline solution and washed and filtered in tap water before being used as feed.

### 4.2 Imaging and microscopy

Individual *H. vulgaris* and *H. magnipapillata* animals placed in a petri dish with a drop of ‘M’ solution to prevent desiccation and imaged using a Leica dissection microscope S8 APO illuminated with a Leica (24 DC) LED control unit and equipped with an EC 3 camera (Leica Microsystems, Germany). The onset of turbulence was recorded using autofocus and automatic exposure settings in video mode on a Canon EOS 1000D camera (Canon Inc., Japan).

### 4.3 Flow chamber and detaching *Hydra*

A syringe pump (PhD Ultra, Harvard Apparatus, USA) with a 20 ml plastic syringe (BD Biosciences, India) was connected to polycarbonate tubing of inner diameter (I.D.) 3 mm (BioRad, USA) and further connected with an adaptor (BioRad, USA) to a tube of I.D. 0.8 mm, taped to the bottom of a glass trough (Figure 2(a)). For *Hydra* detachment experiments, the animals were allowed to attach to a glass coverslip (MicroAid, Pune, India) such that the animal was at a distance of 0.5 cm from the tube outlet, in line with the fluid flow. Flow experiments typically involved increasing the flow rate from 10 ml/min with increments of ∼ 2 ml/min until the animal detached due to the force generated. Those experiments in which the *Hydra* was either attached to the substrate with its tentacles, or did not attach at all, or failed to detach at all flow rates were ignored in analysis of detachment shear stress.

### 4.4 Modulating substrate stiffness

For experiments to measure the effect of substrate stiffness, a 0.75 mm thick 5% polyacrlyamide gel was prepared using a 5 ml solution of 5% acrylamide and 0.22% bisacrylamide, 25 *μl* APS (1/200 volume) and 2.5 *μl* TEMED (1/2000 volume) (all reagents Sigma-Aldrich, Mumbai, India) and curing for 15 min between two plates layered with water in a standard polyacrylamide gel electrophoresis (PAGE) setup (BioRad, USA). This gel of stiffness ∼ 8 kPa (Tse and Engler, 2010), was layered on the coverslip and the *Hydra* was allowed to attach to the gel. The remainder of the experiment was performed in a manner similar to the experiments for detaching *Hydra* from glass coverslips.

### 4.5 Data analysis

Images of *Hydra* and the onset of turbulence were processed using ImageJ (Schneider et al., 2012). Fitting data to functions was performed using the non-linear fitting tool (*nlinfit*) in MATLAB (Mathworks Inc., MA, USA), as was all plotting. Statistical testing of mean shear stresses was performed by comparing the means by a Student’s t-test.

## Acknowledgements

We would like to acknowledge the kind gift of *Hydra magnipapillata* and *Hydra vulgaris* by Sanjeev Galande and help from Girish Ratnaparkhi and Surendra Ghaskadbi in culturing. This work was funded by IISER Pune core funding.

## References

Amimoto, Y., Kodama, R., and Kobayakawa, Y. (2006). Foot formation in Hydra: a novel gene, anklet, is involved in basal disk formation. Mech. Dev., 123(5):352–361.

Annandale, N. (1911). Freshwater Sponges, Hydrozoids and Polyzoa. Taylor and Francis, Red Lion Court, Fleet Street, London.

Bell, E. C.Gosline, J. M. (1996). Mechanical Design of Mussel Byssus: Material Yield Enhances Attachment Strenght. J. Exp. Zool., 199:1005–1017.

Bode, H., Dunne, J., Heimfeld, S., Huang, L., Javois, L., Koizumi, O., Wester-field, J., and Yaross, M. (1986). Transdifferentiation Occurs Continuously in Adult Hydra. Curr. Top Dev. Biol., 20:257–280.

Brien, P. (1960). The fresh-water hydra. American Scientist, 48(4):pp. 348A, 461–475.

Chen, S. and Springer, T.A. (1999). An automatic braking system that stabilizes leukocyte rolling by an increase in selectin bond number with shear. J. Cell Biol., 144(1):185–200.

Denny, M. W. (1987). Lift as a mechanism of patch initiation in mussel beds. J. Exp. Mar. Biol. Ecol., 113:231–245.

Fujisawa, T. (2006). Hydra is joining the bandwagon. BioEssays, 28(5):560–562.

Gorb, S. N. (2008). Biological attachment devices: exploring nature's diversity for biomimetics. Philosophical Transactions of the Royal Society A: Mathematical, Physical and Engineering Sciences, 366(1870):1557–1574.

Koehl, M. A. R. (1977). Effects of Sea Anemones on the Flow Forces they Encounter. J. exp. Biol., pages 85–105.

Lee, H., Scherer, N. F., and Messersmith, P. B. (2006). Single-molecule mechanics of mussel adhesion. Proc. Natl. Acad. Sci. U.S.A., 103(35):12999–13003.

Lin, Q., Gourdon, D., Sun, C., Holten-Andersen, N., Anderson, T. H., Waite, J. H., and Israelachvili, J. N. (2007). Adhesion mechanisms of the mussel foot proteins mfp-1 and mfp-3. Proceedings of the National Academy of Sciences of the United States of America, 104(10):3782–6.

Neal, A., Simoes, F., and Yule, A. (1996). Interactions between shear rates and biofilms affecting exploratory behaviour by cyprids ofelminius modestus (cirripedia). Marine Biology, 127(2):241–246.

Reddy, C. P., Barve, A., and Ghaskadbi, S. (2011). Description and phylogenetic characterization of common hydra from India. Curr. Sci., 101(6):736–738.

Rodrigues, M., Leclère, P., Flammang, P., Hess, M. W., Salvenmoser, W., Hobmayer, B., and Ladurner, P. (2016a). The cellular basis of bioadhesion of the freshwater polyp hydra. BMC Zoology, 1(1):3.

Rodrigues, M., Ostermann, T., Kremeser, L., Lindner, H., Beisel, C., Berezikov, E., Hobmayer, B., and Ladurner, P. (2016b). Profiling of adhesive-related genes in the freshwater cnidarian Hydra magnipapillata by transcriptomics and proteomics. Biofouling, 32(9):1115–1129.

Schneider, C., Rasband, W., and Eliceiri, K. (2012). NIH Image to ImageJ: 25 years of image analysis. Nature Methods, 9:671–675.

Schultz, M. P., Finlay, J. A., Callow, M. E., and Callow, J. A. (2000). A turbulent channel flow apparatus for the determination of the adhesion strength of microfouling organisms. Biofouling, 15(4):243–251.

Shimizu, H., Takaku, Y., Zhang, X., and Fujisawa, T. (2007). The aboral pore of hydra: evidence that the digestive tract of hydra is a tube not a sac. Development genes and evolution, 217(8):563–568.

Stoecker, H. (2004). Taschenbuch der Physik. Verlag Harri Deutsch, 5 edition.

Sugiyama, T. and Fujisawa, T. (1977). Genetic Analysis of Developmental Mechanisms in Hydra I. Sexual Reproduction of Hydra Magnipapillata and Isolation of Mutants. Develop., Growth and Differ., 19(3):187–200.

Touchette, B. W., Marcus, S. E., and Adams, E. C. (2014). Bulk elastic moduli and solute potentials in leaves of freshwater, coastal and marine hydrophytes. Are marine plants more rigid? AoB Plants, 6.

Tse, J. R. and Engler, A. J. (2010). Preparation of hydrogel substrates with tunable mechanical properties. In Curr. Prot. Cell Biol., chapter 10. Wiley.

Vogel, S. and LaBarbera, M. (1978). Simple Flow Tanks for Research and Teaching. BioScience, 28(10):638–643.

Wagner, G. (1905). Memoirs: On Some Movements and Reactions of Hydra. Q. J. Microsc. Sci., s2(48):585–622.

Waite, H. J. (2002). Adhesion a la Moule. Integr. Comp. Biol., 42:1172–1180.

Watanabe, H., Hoang, V., Maẗtner, R., and Holstein, T. (2009). Immortality and the base of multicellular life: Lessons from cnidarian stem cells. Semin. Cell Dev. Biol., 20(9):1114–1125.

